# *In vivo* imaging of myonuclei during spontaneous muscle contraction reveals non-uniform nuclear mechanical dynamics in *Nesprin/klar* mutants

**DOI:** 10.1101/643015

**Authors:** Dana Lorber, Ron Rotkopf, Talila Volk

## Abstract

Muscle contractions produce reiterated cytoplasmic mechanical variations, which impact the nuclear membrane and potentially influence nuclear mechanotransduction. It is unclear, however, whether the mechanical dynamics of individual myonuclei changes during each contractile wave, and whether mutants whose connection to the cytoskeleton is impaired are subjected to different mechanical input.

To monitor nuclear mechanical dynamics *in vivo*, we imaged myonuclei along muscles during multiple spontaneous muscle contractile events, within live and intact *Drosophila* wild-type or *Nesprin/klar* mutant larvae. The data were subsequently analyzed and quantified aiming to reveal potential changes in the mechanical parameters of nuclear dynamics during muscle contraction in the *Nesprin/klar* mutant. Our results show that all myonuclei in control larvae exhibited comparable dynamics in the course of multiple contractile events. In contrast, myonuclei of homozygous mutant larvae lacking the Nesprin-like gene *klar* displayed differential dynamics relative to wild type, and higher variability between myonuclei at distinct positions along individual myofibers. Estimation of the drag force applied on individual myonuclei revealed that force fluctuations in time were considerably higher in the *Nesprin/klar* mutant myonuclei relative to control, reflecting a significant variability in the mechanical dynamics of individual myonuclei during contractile waves. The variable mechanical dynamics along the muscle fiber and the higher variance of *Nesprin/klar* mutant myonuclei may lead to altered nuclear mechanotransduction. Since mutations in *Nesprin* genes lead to devastating muscle and cardiac human diseases, our findings may provide new insight into the mechanism underlying these pathologies.

## Introduction

Muscles are particularly receptive to mechanical stimulation and respond to increased mechanical load by changing transcriptional and translational levels (Egan and Zierath, 2013; Hawley et al., 2014). Recent evidence suggests that the nuclear envelope transmits mechanical cues into the nucleus, which are essential to control epigenetic state and gene expression (Fedorchak et al., 2014; Martins et al., 2012). However, the way such mechanical signals spread over hundreds of myonuclei comprising individual myofibers remains to be elucidated (Hector et al., 2015; Rindom and Vissing, 2016). To reveal the mechanical dynamics of myonuclei during muscle contraction it is essential to accumulate quantitative information during the course of contractile events. Obtaining such information in an intact live organism is challenging due to the difficulty to follow the path of individual myonuclei within specific myofibers in a single contractile event.

Cytoplasmic mechanical signals are transmitted to the nucleus mainly via the Linker of Nucleoskeleton and Cytoskeleton (LINC) complex (Lombardi and Lammerding, 2015; Rothballer and Kutay, 2013). This evolutionarily conserved complex (Razafsky 2014, Mellad et al., 2011) is comprised of cytoskeletal-associated Nesprins, proteins that contain **K**larsicht, **A**NC-1, **S**yne **H**omology (KASH) domain. Nesprins directly bind to **S**ad1p, **UN**C-84 (SUN) domain proteins at the perinuclear space, thereby forming a link between the cytoskeleton and the nucleoskeleton (Cartwright and Karakesisoglou, 2014; Mellad et al., 2011). Previous studies indicated the essential contribution of the LINC complex to nuclear morphology (Banerjee et al., 2014; Rothballer and Kutay, 2013), nuclear migration and positioning (Bone et al., 2014; Rajgor and Shanahan, 2013; Stroud et al., 2017), regulation of nuclear lamina proteins, and chromatin organization (Mellad et al., 2011a; Osmanagic-Myers et al., 2015; Rothballer and Kutay, 2013).

Vertebrate muscles express multiple LINC complex genes, impeding functional analysis of the complex. The *Drosophila* genome contains only two KASH encoding genes, namely *klarsicht* (*klar*) (Xie and Fischer, 2008) and *Msp300* (Elhanany-Tamir et al., 2012; Xie and Fischer, 2008; Zhang et al., 2002)), and a single SUN encoding gene, *klaroid* (koi) (Kracklauer et al., 2007), which facilitates dissecting their functional contribution. Previous analyses indicated that in muscles, *Msp300/Nesprin* is essential for nuclear positioning through mediating association of the nuclear membrane with *Drosophila* Titin (DTitin) at the Z-discs (Elhanany-Tamir et al., 2012). Furthermore, Klar and Msp300 mediate connection of the microtubule network with the nuclear membrane (Elhanany-Tamir et al., 2012; Wang et al., 2015). In addition to nuclear position defects, LINC complex mutant myonuclei exhibited differential nuclear size, variable DNA content, and dysregulated DNA replication (endoreplication). The variability in DNA content in myonuclei of single myofibers implied a novel function of the LINC complex, namely in equalizing the degree of DNA endoreplication among numerous myonuclei of individual myofibers (Wang et al., 2018).

Despite the wide tissue distribution of the LINC complex, mutations in the genes coding for its various components lead primarily to defects in force generating tissues such as skeletal muscles and cardiomyocytes, implying its unique functional contribution in these tissues. To study myonuclear dynamics during muscle contraction, we developed a device that allows imaging of larval body wall muscles in live intact larvae. Unlike other experimental systems, our approach does not require muscle isolation (Barash et al., 2005; Drost et al., 2003; Eldred et al., 2010; Hakim et al., 2013), use of anesthetized organism (Sloboda et al., 2013; Zhang et al., 2010), or complete immobilization of the model (Csordás et al., 2014; Ghannad-Rezaie et al., 2012; Mondal et al., 2012; Zhang et al., 2016), and thus allows to record and estimate nuclear dynamics in intact live organisms.

Significantly, we found substantial differences in the spatiotemporal dynamics of *Nesprin/klar* mutant myonuclei relative to control along the muscle fiber during muscle contraction. Our results suggest that in contrast to control muscles, where synchronized myonuclear dynamics along the entire muscle fiber is maintained in the course of muscle contraction, Nesprin/klar mutant myonuclei lack such synchronization and uniformity. We assume that myonuclear dynamics during muscle contraction correlates with the net forces acting on the nucleus, derived mainly from viscous drag force (Forgacs and Newman, 2005). Thus, the differences between the dynamics of the control and mutant nuclei during contraction may result with different forces applied on the myonuclei during contraction.

## Materials and Methods

### The minimal constraint device for imaging live intact Drosophila larvae

The center of the device is a thin plastic bar positioned in a larger frame, with a groove crossing the frame and bar from side to side, as shown in Figure 1A. The larva is placed in the groove that allows free body movement but prevents bending. Glass capillaries, which are aligned with the larval body, are attached to the head and tail to prevent it from crawling. A coverslip glass covers the larva and glass capillaries, which are flipped together before the device is placed on the microscope stage. The larvae are inserted into the groove in the central bar with forceps and then are rotated with a fine brush along its axis, such that the longitudinal muscles will face the coverslip glass. The head and tail are glued (Superglue liquid, Pattex, Australia) to a glass capillary (Drummond Scientific Company, PA, USA, cat. #1-000-0010, O.D 0.026”, I. D. 0.0079”), as shown in Figure 1B. Next, the frame with the bar and larva is inserted into the device base. The bar is filled with hydrogel that damps the vertical movement of the larva, while it is still soft enough to enable motility. In addition, the hydrogel keeps the larvae in a moist environment. The hydrogel used was alginate, a common material in the biomedical and food industries (Olatunji, 2015; Wan et al., 2009), which polymerizes at room temperature in the presence of bivalent cations. A solution of 4% alginate (cat. # 180947, Aldrich) dissolved in 0.9% saline (NaCl, cat. # 0277, J. T. Baker) was polymerized with a solution of 0.8 M of CaCl_2_·2H_2_O (cat. # 2382.0.0500, Merck).

**Figure 1:**
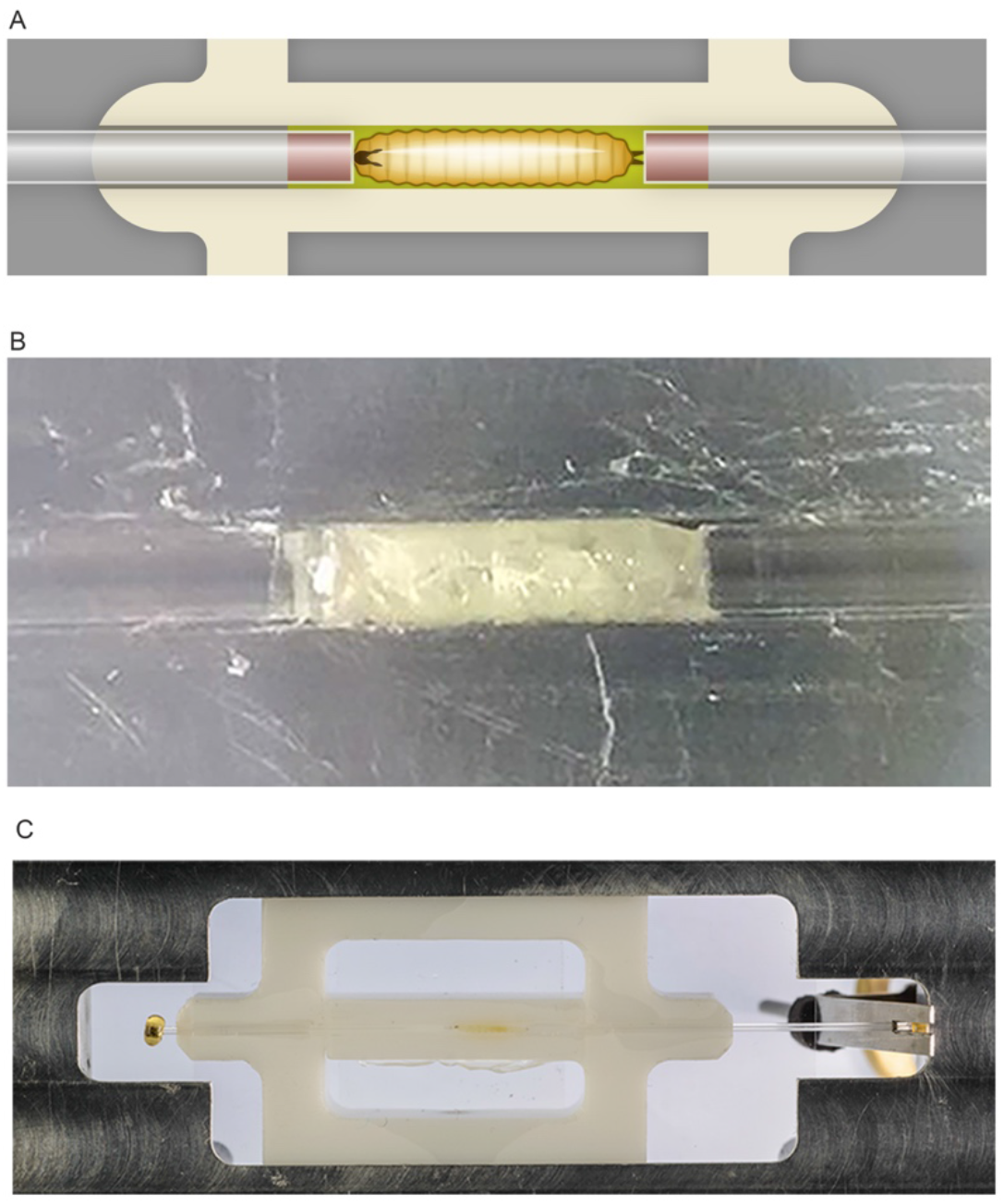
The minimal constraint device for imaging live intact *Drosophila* larvae. (A) Configuration of the device: The larva is placed along a groove with two glass capillaries filled with glue attached to its head and tail. The space between the larva and plastic bar is filled with hydrogel. (B) An image of a larva in the groove attached to the capillaries. (C) The plastic bar containing a larva is assembled into the base of the device.

#### Flies

The flies used in this study, P{UAS-mCherry.NLS}2 and P{GAL4-Mef2.R}2, were obtained from Bloomington Stock Centre. The flies were recombined on the second chromosome and crossed with flies with endogenously EGFP-tagged *sallimus* (obtained from Belinda Bullard, Department of Biology, University of York). For the analysis of *klar* homozygous larvae, the *klar^1-18^/TM6,Tb* allele, which lacks all *klar* coding exons (obtained from Michael Welte, Department of Biology, University of Rochester) was recombined with the Sls-GFP insertion, and the recombined line P{UAS-mCherry.NLS}2, P{GAL4-Mef2.R}2 was combined with *klar^1-18^/TM6,Tb*.

#### Imaging

All the experiments were performed with a VisiScope CSU-W1 Spinning Disk Confocal Microscope (Visitron Systems, Puchheim, Germany) based on an Olympus IX83 inverted microscope, a Yokogawa CSU-W1 scan head, and a PCO Edge4.2 sCMOS camera. The images were acquired with a 20X air objective (NA=0.75), which enables visualization of an entire intact longitudinal muscle during its activity.

Laser lines of 561 nm and 488 nm were used to excite the mCherry and GFP fluorescent markers, respectively, and the exposure time of each image was 20 ms. In order to shorten the time between acquisitions of consecutive images, an emission dual band filter (cat. # 59022m, Chroma Technology Corp, USA) was added to the microscope, which isolates the emitted light to two separate monochromatic wave lengths. The merged images of the myonuclei and sarcomeres were used to find the edges of the muscles and to determine which myonuclei belong to the tested muscle. Spontaneous muscle contractions were recorded continuously, up to 500 pairs of images in a single sampling, which is the limitation of the sampling system.

For quantification, only full muscle contractions were considered. Contraction onset and completion were determined when at least in two consecutive images the nuclei did not change their position relative to each other.

#### Quantification of myonuclear dynamics

To determine the spatiotemporal dynamics of each nucleus along the muscle fiber during a contractile event, its displacement in time was measured. The location of each nucleus in each image, throughout a single contractile event, was measured using the manual tracking plug-in from ImageJ. The displacement d of a nucleus that occurred between two successive sampling times t_1_ and t_2_ was determined as follows:

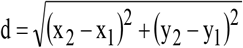

where (x_1_, y_1_) is the location of the middle of the nucleus at t_1_ and (x_2_, y_2_) is the location at t_2_, as shown in Figure 2A.

**Figure 2:**
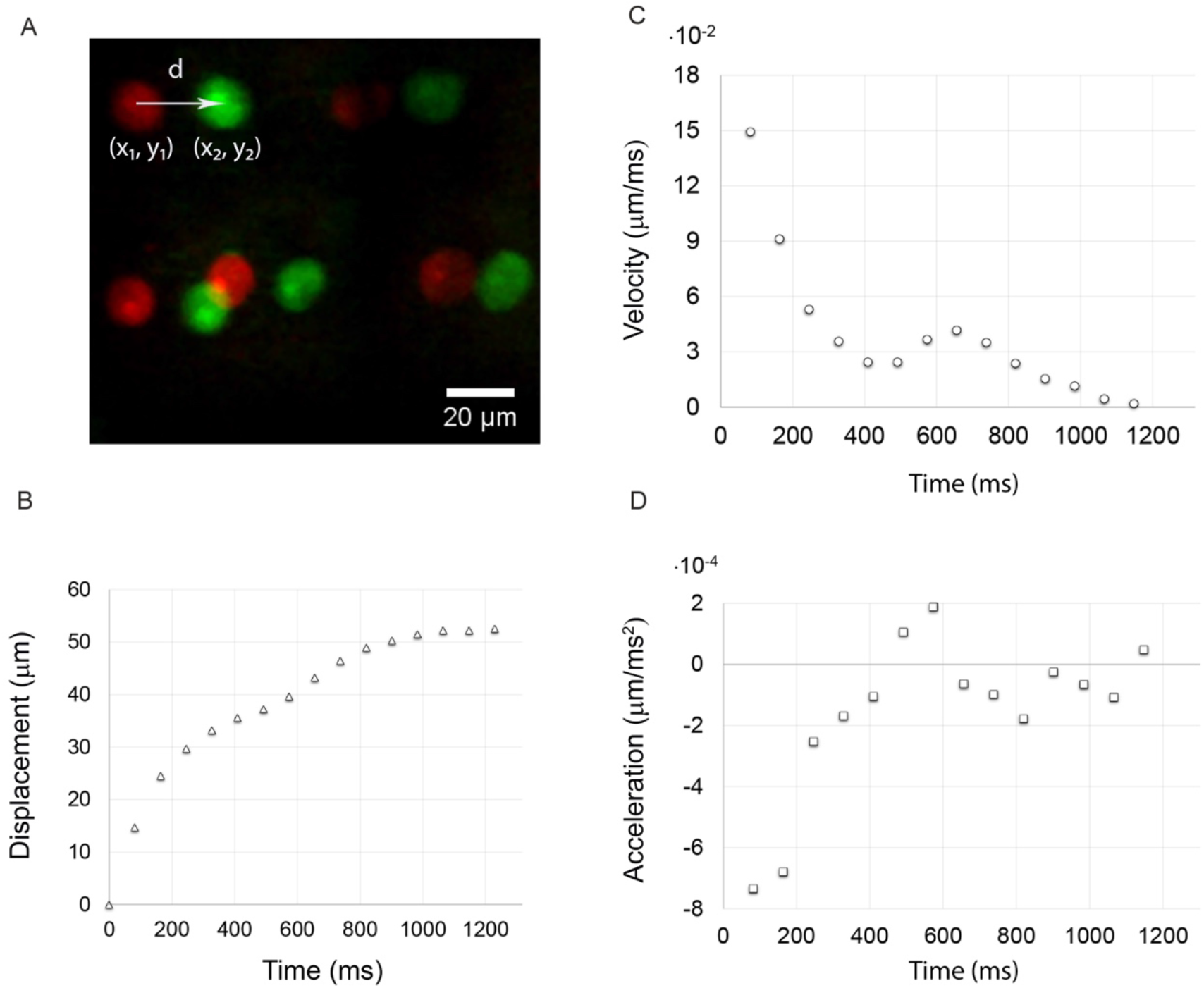
Measurements of nuclear displacement and parameters calculations. An example of two sequential images of contractile muscle demonstrating myonuclear displacement of 5 nuclei during a time interval of 82 ms (red is t=0 and green is t=82 ms). (B) The displacement in time of a single nucleus during a complete contraction. (C) The velocity of a nucleus during contraction was calculated based on the temporal displacement function. (D) The acceleration of a nucleus during contraction was calculated based on the displacement function.

At the end of each contraction, the muscle returned to its original position. Hence, the displacement measurements relate to a stationary frame of reference and there is no need to correct for relative movement of the larva and device.

The sampling times that were obtained from the sampling system enabled to derive the displacement function in time. To calculate the velocity in time v(t) and acceleration in time a(t), the first and second discrete numerical central derivatives with unevenly spaced points were calculated, respectively, based on Taylor’s expansion (Gupta, 2015). This approach was used because the sampling rate of the microscope was found to be inconsistent. Figure 2B-D shows a representative example of the three functions in time, as calculated for a single nucleus in a single contraction. Based on these three functions in time, several physical parameters were calculated, such as average velocity and average acceleration during contraction. We also calculated the path, which is the trajectory length of a nucleus during contraction, and the net displacement, which is the distance from the starting location to the end one.

For low Reynolds’ number (Re<1), the viscous drag force F on a spherical body of radius R moving in the fluid can be calculated using Stokes law: F=6πηRv, where η is the viscosity of the fluid and v is its velocity. (Fuks NA, 1964). The force time derivative dF/dt, assuming the viscosity of the fluid and the geometry of the body are constant, yields:

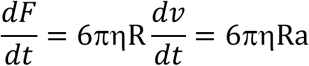

where a is the acceleration of the sphere.

To calculate the drag force applied on an oblate axisymmetric ellipsoid moving in a fluid, a geometric correction factor k is introduced to Stokes law: F=6πηrvk, where r is the equatorial radius of the ellipsoid. k is dependent on the ratio β, between r and other axis of ellipsoid b, and on the direction of the flow in relation to the axis of revolution. When the flow is perpendicular to b (Fuks NA, 1964), then:

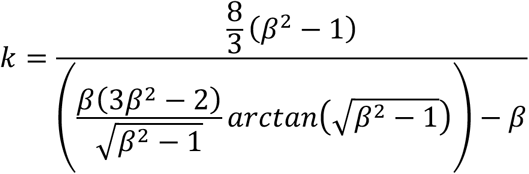

To estimate the average height of control and *klar* mutant nuclei, in another set of experiments, larvae were tilted to allow visualization of the nuclear and muscle sagittal plane, enabling imaging of the nuclear height at high resolution. Using the modified Stokes equation allows to estimate the average force (AF) and average force time derivative (AFTD) applied on the myonuclei during contraction.

#### Programming

A Perl script was used to read the values from the Excel worksheet and then to perform the required functions. The output was written to a new worksheet.

#### Statistics

When one average value was used per larva, differences between WT and mutant larvae were tested using *t*-tests. Differences accounting for location (anterior vs. middle vs. posterior) were analyzed using a repeated-measures ANOVA, accounting for the random effect of each larva, i.e. one model including the group effect (control vs. mutant), and two models for each group separately. Contrasts within a certain location were tested using *t*-tests. Variance comparisons were done using *F*-tests. All statistical analyses were done using R v. 3.5.1.

### Results

#### The minimal constraint device for imaging live intact Drosophila larvae

To record myonuclear dynamics during muscle contractile events, we developed a device that enables tracking myonuclei in muscles of intact, live 3^rd^ instar *Drosophila* larvae under a spinning disc microscope (Figure 1A-C, and see Materials and Methods). The myonuclei were labelled with mCherry fused to a nuclear localization signal (mCherry-NLS) driven by *mef2-GAL4* driver, and sarcomeres were visualized by endogenous GFP insertion into the *D-Titin* gene (*sallimus-GFP*). Longitudinal muscles were routinely recorded using a 20X objective, capturing a length of 665.6 μm, enabling image acquisition of an entire myofiber from anterior to posterior segmental borders.

To evaluate the dynamic behaviour of the myonuclei in the course of a single contractile event, we calculated the displacement in time of each nucleus based on the change in its x-y position from frame to frame. This was followed by calculating the velocity and acceleration for each of the myonuclei in numerous movies taken. A representative example for the displacement of a few myonuclei that occurred between two consecutive images is shown in Figure 2A, in which the colour of myonuclei was artificially changed from red (earlier point) to green (later point, t=82 ms). Note that the displacements (d) of these myonuclei are not uniform during this time interval. A representative example of the displacement of a single myonucleus in time during full contraction is shown in Figure 2B, and its velocity and acceleration values are shown in Figure 2C and 2D, respectively. Note that in this example, nuclear displacement increased rapidly in the first 800 ms and then tapered off. Nuclear velocity decreased with time, and the absolute values of nuclear acceleration varied in time.

Representative single still images of relaxed or fully contractile myofibers of control larvae, which were extracted from Supplementary Movie 1, are shown in Figure 3A-B’. To follow nuclear dynamics in a LINC complex representative mutant, we used *Nesprin/klar* homozygous larvae in combination with the fluorescent markers described above. Representative single confocal still images of relaxed or fully contractile myofibers of *Nesprin/klar* homozygous mutant muscles, which were extracted from Supplementary Movie 2, are shown in Figure 3C-D’. Consistent with previous studies, the *Nesprin/klar* mutant muscles displayed uneven distribution of the myonuclei and, in addition, exhibited variable size (Figure 3C-D’). The trajectories of each of the nuclei shown in Figure 3A-D’ are diagrammed in Figure 3E,F (E for control and F for *Nesprin/klar* mutant muscles). Notably, differential displacement of myonuclei was observed in control and *Nesprin/klar* mutant muscles. These results suggest that the dynamics of the *klar* mutant myonuclei differs from that of control.

**Figure 3:**
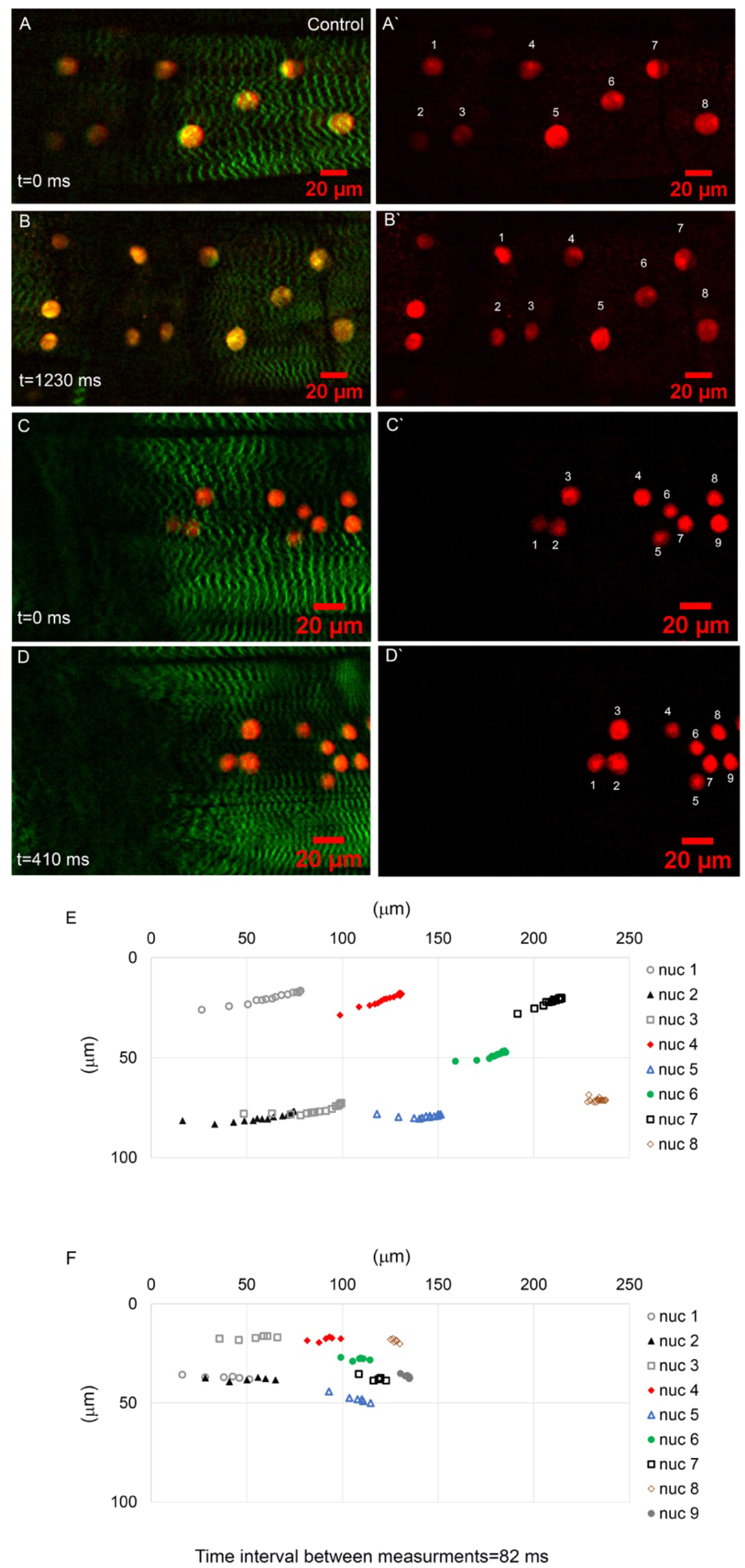
Detection of myonuclei during contraction. (A-D) control (A,B,A’B’) and *Nesprin/klar* mutant (C,D,C’D’) myonuclei labeled with Cherry-NLS driven by *Mef2-GAL4* in relaxed (A,A’,C,C’) or fully contractile (B,B’,D,D’) muscles. Sarcomeres are labeled with Sls-GFP. Scale bars are 20 μm. (E) The path that each of the control myonuclei shown in A-B’ undertakes during a full contractile event. (F) The path that each of the *Nesprin/klar* myonuclei shown in C-D’ undertakes during a full contractile event.

#### Quantitative analysis of myonuclear dynamics of control and Nesprin/klar mutant muscles

To reveal whether nuclear dynamics during muscle contraction of *Nesprin/klar* mutant myonuclei differs from that of control, we calculated the time dependent displacement, velocity, and acceleration of myonuclei in numerous contractile events in control and *Nesprin/klar* mutant larvae. In both groups (control or *Nesprin/klar* mutant), measurements of nuclear displacements were taken from five 3^rd^ instar larvae, in which up to 10 events of muscle contractions were quantified from a single muscle, and for all visible myonuclei. Table 1 summarizes the number of myonuclei analysed for each muscle, and the number of contractile events analysed for each group.

**Table 1:** Summary of the number of nuclei analysed

| Exp. | Num. of nuclei | Num. of contractions | Exp. | Num. of nuclei | Num. of contractions |
| --- | --- | --- | --- | --- | --- |
| Control 1 | 15 | 6 | *klar* 1 | 6 | 7 |
| Control 2 | 9 | 8 | *klar* 2 | 17 | 7 |
| Control 3 | 11 | 7 | *klar* 3 | 7 | 6 |
| Control 4 | 12 | 7 | *klar* 4 | 9 | 10 |
| Control 5 | 13 | 10 | *klar* 5 | 13 | 8 |
| sum | 60 | 38 | sum | 52 | 38 |

The quantified mechanical parameters included contraction time (Figure 4A), path (Figure 4B), average velocity (Figure 4C and 4C’), and average acceleration (Figure 4D and 4D’).

**Figure 4:**
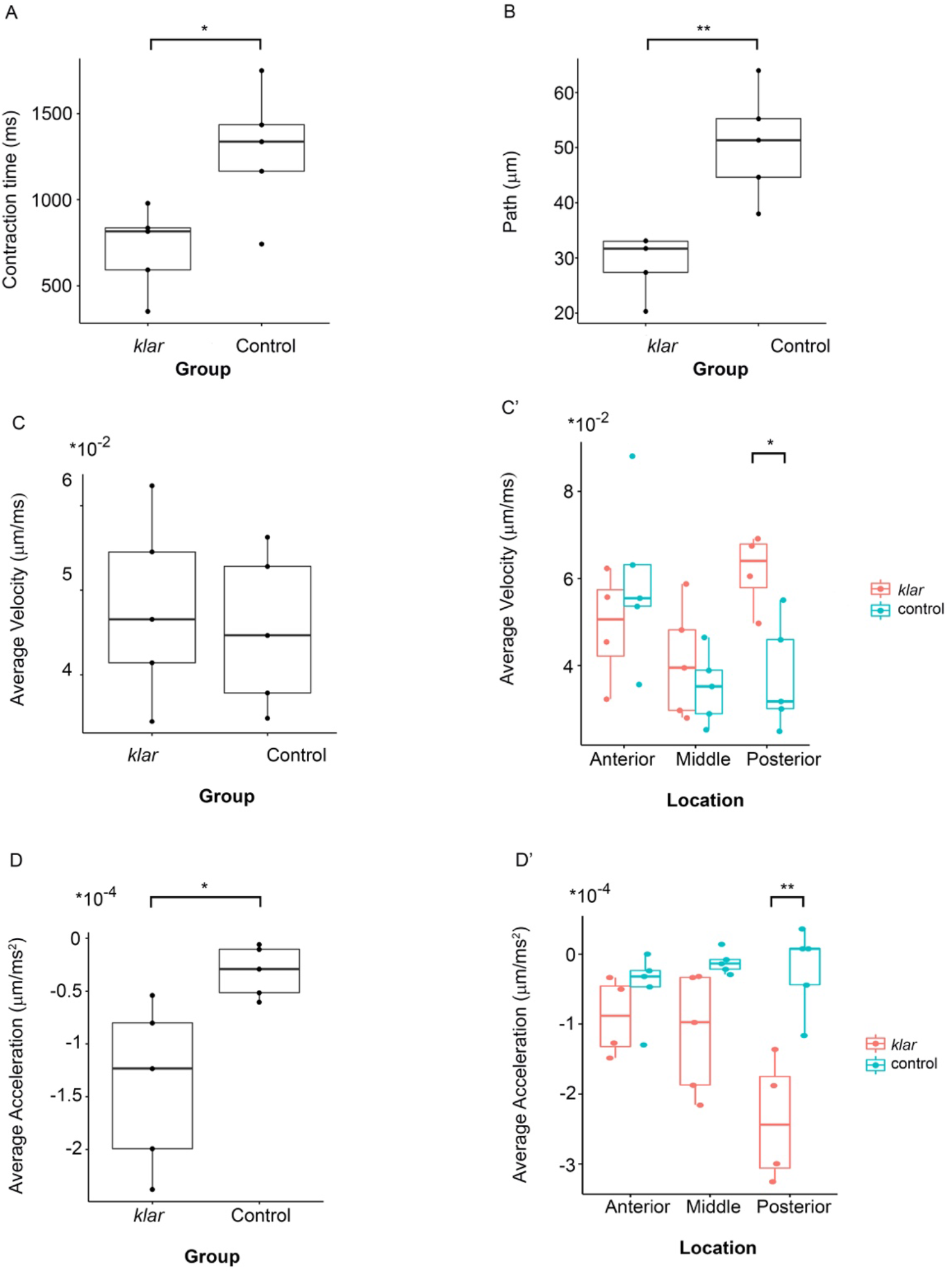
Quantification of nuclear mechanical parameters in control and *Nesprin/klar* larvae during muscle contractile waves. The average contractile time (ms) in *klar* and control myonuclei. Each point represents the average of 6 to 10 contractile events per larvae in all panels (as detailed in Figure 3). The averaged contraction time for control was 1286.32 ± 371.55 ms and for *klar* mutant 715.16 ± 245.68 ms (*t*-test, *p*=0.0024). (B) The average path during a single contractile event of all the nuclei measured per larvae. The average path was 50.64 ± 9.93 μm for control and 29.1 ± 5.43 μm for *klar* mutants (*t*-test, *p*=0.005). In each muscle there were 6 to 17 nuclei, thus each point represents 42 to 130 measurements. (C) The average velocity of myonuclei per individual contractile event. No significant difference was observed. (C’) The average velocity of anterior, middle, or posterior myonuclei. A significant difference between the average velocity of posterior myonuclei between *klar* mutants and control larvae was observed (*p*=0.012). (D) The average acceleration values of myonuclei in *klar* mutants (−1.39·10^−4^±7.8·10^−5^ μm/msec^2^) and control larvae (3.14·10^−5^±2.42·10^−5^ μm/msec^2^) (*p*=0.034). The variance of the average acceleration was significantly higher in *klar* mutants versus control (*p*=0.044). (D’) Average acceleration values calculated per anterior, middle, or posterior myonuclei. A significant difference was observed between control and *klar* mutant myonuclei at the posterior position (*p*=0.009).

The average duration of a single contractile event in control larvae was significantly longer compared to that of *Nesprin/klar* mutant muscle (1286.32 ± 371.55 ms compared to 715.16 ± 245.68 ms, *t*-test, *p*=0.024). Then, the path that each myonucleus traversed during a single contractile event was measured by the summation of all displacements that occurred between consecutive sampling points during contraction. Each point in Figure 4B represents the average path of all the myonuclei from one muscle of a single larva, calculated from 6 to 17 myonuclei that underwent 6 to 10 contractile events, so that each point represents the average of corresponding 42 to 130 different values. The results indicated that the average path traversed by control myonuclei in an individual contractile event was significantly larger relative to that of *Nesprin/klar* myonuclei (50.64 ± 9.93 μm relative to 29.1 ± 5.43 μm, *t*-test, *p*=0.005). A similar tendency was observed when comparing the displacement of the myonuclei between control and *Nesprin/klar* mutant muscles (not shown). Taken together, these results indicate a significant difference between the extent of muscle contraction in *Nesprin/klar* versus control muscles.

Next, the average velocity during contraction was calculated for control and *Nesprin/klar* myonuclei per larva (Figure 4C). No significant difference between control and *Nesprin/klar* mutant groups was observed (*t*-test, *p*=0.7). Further analysis addressed nuclear dynamics at distinct locations along myofibers. The two myonuclei closest to the anterior segmental border were defined as “anterior” myonuclei, those closest to the posterior segmental border were defined as “posterior” myonuclei, and the two myonuclei at the geometric center of the myofiber were defined as “middle” myonuclei. Figure 4 C’ shows that the average velocity of control myonuclei was uniform at distinct positions along the fiber (*p*=0.065). In contrast, it was significantly variable in *Nesprin/klar* mutant myofibers (*p*=0.032) (Figure 4C’), primarily because the average velocity of the posterior myonuclei significantly differed from myonuclei at the anterior and middle positions. Notably, the average velocity of posterior *Nesprin/klar* mutant myonuclei was higher than that of corresponding control myonuclei (*p*=0.012) (Figure 4C’).

Further calculation of the acceleration values of all the myonuclei during all the contractions indicated that *Nesprin/klar* mutant myonuclei exhibited higher absolute average acceleration values relative to control (*p*=0.034) (Figure 4D). In control myonuclei, the average acceleration was −3.14·10^−5^±2.42·10^−5^ μm/msec^2^, and in *Nesprin/klar* myonuclei it was −1.39·10^−4^±7.8·10^−5^ μm/msec^2^, an order of magnitude higher. Note that in this case, negative acceleration values represent a decrease in velocity rather than direction of movement. Notably, the variance of the average acceleration values was significantly higher in the *Nesprin/klar* mutant muscles (*p*=0.044). Moreover, whereas the average myonuclear acceleration during contraction in control myofibers was uniform at distinct positions along the myofibers (*p*=0.39) (Figure 4D’), in *Nesprin/klar* mutant myofibers, this homogeneity was lost and the average myonuclear acceleration values differed, primarily due to an increase in acceleration of the posterior mutant myonuclei (*p*=0.025) (Figure 4D’). A significant difference in the average acceleration values between control and *Nesprin/klar* mutant was also noted at the posterior position (*p*=0.009) (Figure 4D’).

In summary, our analysis of nuclear mechanics during contraction indicates that control myonuclei appear to be exposed to homogenous mechanical dynamics, whereas the *Nesprin/klar* mutant myonuclei experience variable mechanical dynamics, indicated by the differential velocity and acceleration values along the fiber during muscle contraction.

#### Estimation of the average force and its time derivative applied on the myonuclei during contraction

To estimate the average drag force (AF) exerted on the myonuclei during contraction, we used Stokes law (see Materials and Methods). The viscosity value used to calculate the AF was chosen on the basis of results obtained from several studies that measured human muscle viscosity *in vivo* (Chakouch et al., 2016; Debernard et al., 2013; Gennisson et al., 2010; Hoyt et al., 2008). These calculations displayed a range of 0.7 Pa·s to 10.6 Pa·s for relaxed muscles and 11.6 Pa·s to 12.6 Pa·s for different levels of contractions and loads. We chose a value of 10 Pa·s, which reflects the average value of the viscosity between relaxed and contractile muscle *in vivo*.

The equatorial radius was calculated for each nucleus and was equal to half the average of the measured x-y axes. The average height of control nuclei was determined to be 6.96 μm and that of *klar* mutant nuclei was found to be 6.85 μm, a difference which is not statistically significant. Furthermore, no correlation between the length of the nuclei along the muscle fiber and their height was observed.

No significant difference was observed between the AF applied on myonuclei during contraction of control or *Nesprin/klar* myonuclei (Figure 5A), (*t*-test, *p*=0.73). Significantly, however, we observed a substantial difference between the Average Force Time Derivative (AFTD) obtained for myonuclei in control versus *Nesprin/klar* myonuclei (Figure 5B). On average, *Nesprin/klar* mutant myonuclei were found to be subjected to higher absolute values of AFTD relative to control (*p*=0.033), with an order of magnitude higher variance (*p*=0.048). Furthermore, AFTD values differed for myonuclei at different positions along the myofiber in *Nesprin/klar* mutant muscles (*p*=0.00165), whereas control myonuclei showed uniformity along the myofiber (*p*=0.3) (Figure 5B’).

**Figure 5:**
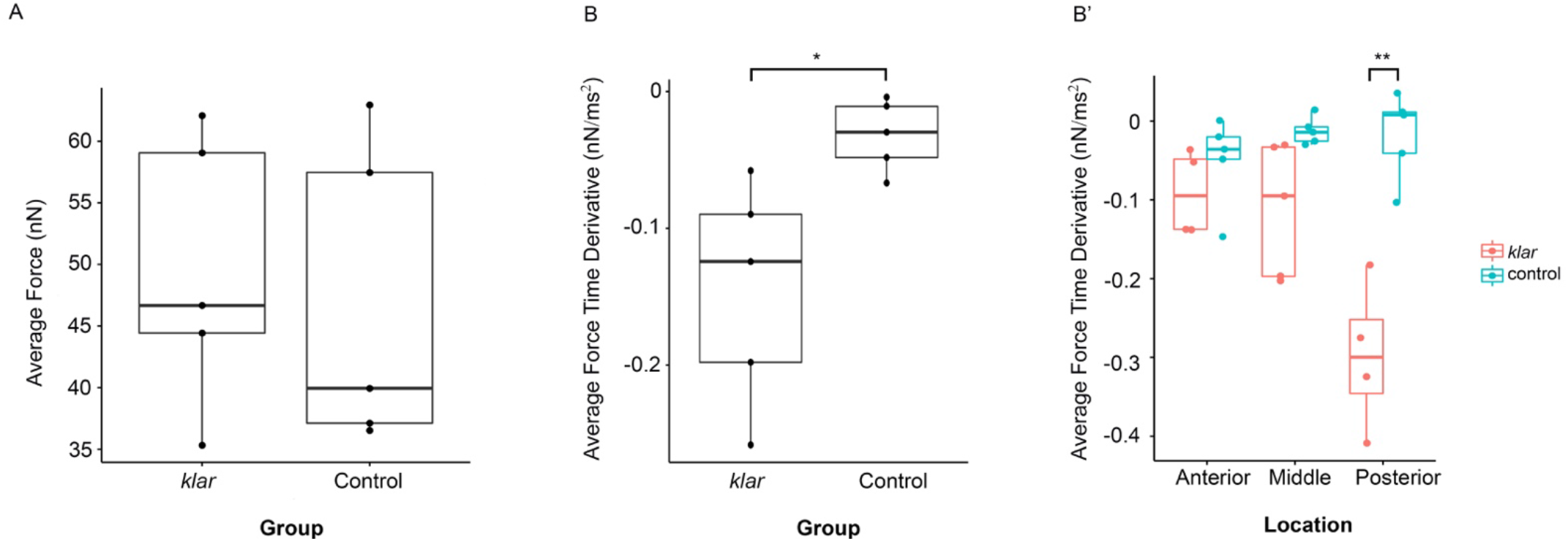
Estimation of the drag force applied on myonuclei during contraction. (A) The average drag force values for control and *Nesprin/klar* mutants do not differ (*p*=0.73). (B) The absolute average force time derivative (AFDT) during contraction is significantly higher in *Nesprin/klar* versus control myonuclei (*p*=0.033) with an order of magnitude higher variance (*p*=0.048). (C) AFTD values differed along the myofiber in *Nesprin/klar* mutant muscles (*p*=0.00165), whereas control myonuclei displayed uniformity along the myofiber (*p*=0.3).

These results are consistent with the idea that Nesprin/klar, shown previously to mediate nuclear association with the microtubule network, as well as nuclear connections with the sarcomeres, is essential for providing uniform mechanical dynamics for myonuclei during muscle contractile waves.

### Discussion

Mechanical signals transduced across the nuclear membrane have been shown to be critical for regulation of the chromatin state, gene expression levels, and nuclear import of transcription factors (Cho et al., 2017; Elosegui-Artola et al., 2017; Fedorchak et al., 2014; Miroshnikova et al., 2017; Uhler and Shivashankar, 2017). Myonuclei are exposed to variable cytoplasmic mechanical inputs in the course of their contractile waves; however, quantitative information regarding the extent of the actual mechanical forces, as well as their spatial and temporal distribution along the elongated myofiber, in a living organism during spontaneous contractions, is still missing. Here, we provide such quantitative mechanical information regarding myonuclear dynamics at a single myofiber resolution in live organism. In addition, the obtained nuclear dynamics data was compared to that of a LINC-representative mutant, *Nesprin*/*klar*. Our approximations allowed us to evaluate the extent of mechanical drag force applied on myonuclei during muscle contraction in control and *Nesprin/klar* mutant larvae.

A key conclusion from these experiments is that *Nesprin/klar*, shown previously to mediate nucleus-cytoskeleton interaction, is essential for equalizing the mechanical dynamics of the myonuclei and attenuates the drag forces applied on the nuclear membrane of each of the multiple myonuclei that consist a single myofiber. We suggest that providing the myonuclei with stable and homogenous mechanical dynamics facilitates their synchronized response to a large variety of physiological and mechanical conditions.

Our study demonstrates that whereas control myonuclei experience comparable average velocity and acceleration values during contraction and, hence, uniform AF and AFTD, the *Nesprin/klar* mutant myonuclei fail to maintain this homogeneity. Lack of synchronization between the myonuclei observed in *Nesprin/klar* mutants is predicted to cause non-homogenous response to mechanical stimulation produced in the course of muscle contraction. Previous reports describe lack of nuclear size uniformity among myonuclei of *Nesprin/klar* mutant myonuclei in *Drosophila*, as well as in humans (Elhanany-Tamir et al., 2012; Puckelwartz et al., 2010; Stroud et al., 2017). Furthermore, we previously reported that *Drosophila* LINC complex-deficient myonuclei contain variable DNA content as a result of unsynchronized DNA replication (Wang et al., 2018). Variable size and DNA content in *LINC* mutant myonuclei correlates, and might be functionally related to the variable mechanical dynamics observed in the mutant myonuclei. The higher sensitivity of posterior myonuclei observed in *Nesprin/klar* mutant muscles might reflect differential mechanical dynamics in the posterior aspects of the myofiber. Such mechanical variations are presumably sequestered by nuclear association with the cytoskeleton via *Nesprin*/*klar*, and possibly other LINC complex components.

Our measurements indicated that the average contractile duration of the *Nesprin/klar* mutant muscles is significantly shorter than that of control muscles. A partial explanation for this may be our previous finding that Troponin C levels decreased at the transcription, mRNA, and protein levels in *Nesprin/klar* mutants (Wang et al, 2018). Aberrant muscle contraction has been also described in human muscular dystrophies associated with mutations in the LINC complex (Janin et al., 2017; Méjat and Misteli, 2010).

The immediate surroundings of myonuclei include a network of microtubules, sarcoplasmic reticulum, multiple mitochondria and sarcomeres. During muscle contraction and sarcomere shortening, the myonuclei move with the sarcomeres. However, the cytoplasm does not move together with the sarcomeres, thus the nuclei are subjected to friction forces with the cytoplasm content. Based on this assumption, it was possible to give an approximation for the drag force applied on the myonuclei during their movement, using Stokes’ law. Interestingly, although the AF values did not differ between control and *Nesprin/klar* mutant myonuclei, the AFTD and its variance during the contractile events were significantly higher in the *Nesprin/klar* myonuclei. Therefore, force fluctuations in time may represent the main difference between control and *Nesprin/klar* mutant muscles. Such force fluctuations might have long term effects on nuclear activity.

We suggest that the *Nesprin/klar* and possibly the entire LINC complex may serve as a mechanical *low pass filter*, which attenuates the mechanical signals produced during multiple contractile events, resulting in a relatively constant myonuclear mechanical input (Figure 6). This model is consistent with our data demonstrating that the average acceleration and, hence, the AFTD during contraction of control myonuclei was an order of magnitude lower than that measured for the *Nesprin/klar* myonuclei. It also explains why the variance of the acceleration and AFTD values in control myonuclei were significantly lower than those of *Nesprin/klar* myonuclei.

**Figure 6:**
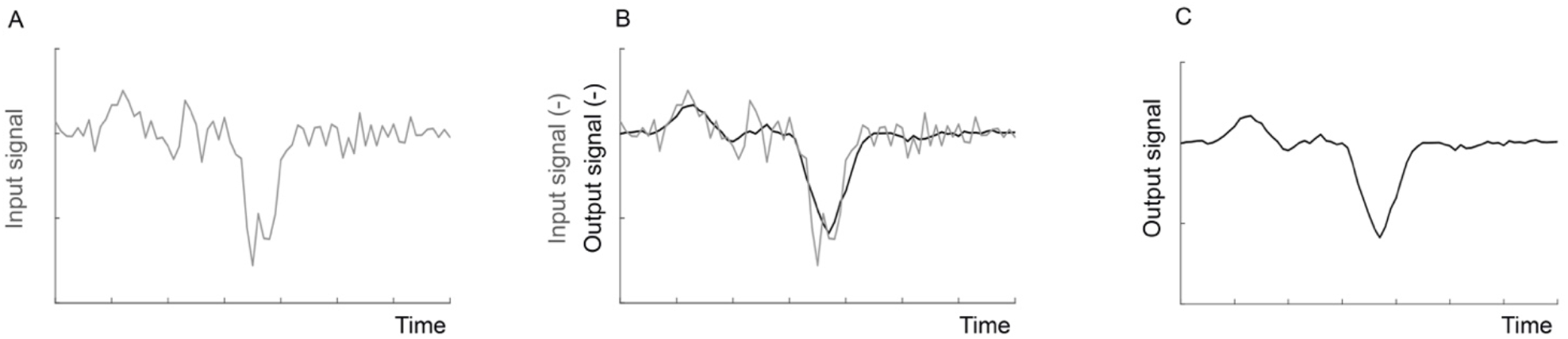
A model for low pass filter. An ideal low pass filter, which can be integrated into mechanical or electrical systems, will eliminate all the frequencies above a designed cut-off from the input signal, and transfer the rest of the input without alternation. (A) The original signal. (B) The original signal, and the output of the same signal after passing through a low pass filter. Note that only fast changes were eliminated. (C) The filtered output signal.

In summary, our results suggest that the LINC complex buffers the nuclear envelope from variable mechanical perturbations that occur during muscle contraction and, thereby, provides a uniform and stable mechanical dynamics for all myonuclei within individual fibres, which is required for homogenous myonuclear response.

## Acknowledgements

We thank the Bloomington Stock Centre for various fly lines, the Developmental Studies Hybridoma Bank (DSHB) for antibodies, and FlyBase for important genomic information. We are grateful for Samuel Safran, from the Department of Chemical and Biological Physics, Weizmann Institute of Science, and his students Dan Deviri and Ohad Cohen for insightful discussions and helpful remarks, Mike Shorkend for writing the codes for our analysis and Nitzan Konstantin for English editing. This study was supported by a grant from the NSF-BSF (BSF application 2016738 to T.V. and NSF application 1715606 to JL) and ISF grant 750/17 (T.V.). FlyBase is supported by a grant from the National Human Genome Research Institute at the U.S. National Institutes of Health #U41 HG000739. Support is also provided by the British Medical Research Council (#MR/N030117/1) and the Indiana Genomics Initiative. The BDSC is supported by a grant from the Office of the Director of the National Institute of Health under Award Number P40OD018537. NIH ICs OD, NIGMS, NICHD and NINDS contribute to funding of this award.

## Author contributions

D.L. conceptualized, investigated, performed formal analysis and wrote the original draft, R.R. performed formal analysis, T.V. supervised, provided funding, reviewed and edited the draft.

## Supplemental data

Movie 1: Representative example of a movie taken from control larvae.

Movie 2: Representative example of a movie taken from *klar* mutant larvae.

## References

Anno T, Sakamoto N, Sato M. 2012. Role of nesprin-1 in nuclear deformation in endothelial cells under static and uniaxial stretching conditions. Biochem Biophys Res Commun 424:94–99. doi:10.1016/j.bbrc.2012.06.073

Banerjee I, Zhang J, Moore-Morris T, Pfeiffer E, Buchholz KS, Liu A, Ouyang K, Stroud MJ, Gerace L, Evans SM, McCulloch A, Chen J. 2014. Targeted Ablation of Nesprin 1 and Nesprin 2 from Murine Myocardium Results in Cardiomyopathy, Altered Nuclear Morphology and Inhibition of the Biomechanical Gene Response. PLoS Genet 10. doi:10.1371/journal.pgen.1004114

Barash IA, Mathew L, Lahey M, Greaser ML, Lieber RL. 2005. Muscle LIM protein plays both structural and functional roles in skeletal muscle. Am J Physiol Cell Physiol 289:C1312–20. doi:10.1152/ajpcell.00117.2005

Bone CR, Tapley EC, Gorjanacz M, Starr DA. 2014. The Caenorhabditis elegans SUN protein UNC-84 interacts with lamin to transfer forces from the cytoplasm to the nucleoskeleton during nuclear migration. Mol Biol Cell 25:2853–2865. doi:10.1091/mbc.E14-05-0971

Brosig M, Ferralli J, Gelman L, Chiquet M, Chiquet-Ehrismann R. 2010. Interfering with the connection between the nucleus and the cytoskeleton affects nuclear rotation, mechanotransduction and myogenesis. Int J Biochem Cell Biol 42:1717–1728. doi:10.1016/j.biocel.2010.07.001

Cartwright S, Karakesisoglou I. 2014. Nesprins in health and disease. Semin Cell Dev Biol 29:169–179. doi:10.1016/j.semcdb.2013.12.010

Chakouch MK, Pouletaut P, Charleux F, Bensamoun SF. 2016. Viscoelastic shear properties of in vivo thigh muscles measured by MR elastography. J Magn Reson Imaging 43:1423–1433. doi:10.1002/jmri.25105

Cho S, Irianto J, Discher DE. 2017. Mechanosensing by the nucleus: From pathways to scaling relationships 1–11.

Csordás G, Varga GIB, Honti V, Jankovics F, Kurucz É, Andó I. 2014. In vivo immunostaining of hemocyte compartments in Drosophila for live imaging. PLoS One 9. doi:10.1371/journal.pone.0098191

Debernard L, Leclerc GE, Robert L, Charleux F, Bensamoun SF. 2013. in Vivo Characterization of the Muscle Viscoelasticity in Passive and Active Conditions Using Multifrequency Mr Elastography. J Musculoskelet Res 16:1350008. doi:10.1142/s0218957713500085

Drost MR, Maenhout M, Willems PJB, Oomens CWJ, Baaijens FPT, Hesselink MKC. 2003. Spatial and temporal heterogeneity of superficial muscle strain during in situ fixed-end contractions. J Biomech 36:1055–1063. doi:10.1016/S0021-9290(02)00461-X

Egan B, Zierath JR. 2013. Exercise metabolism and the molecular regulation of skeletal muscle adaptation. Cell Metab 17:162–184.

Eldred CC, Simeonov DR, Koppes RA, Yang C, Corr DT, Swank DM. 2010. The mechanical properties of drosophila jump muscle expressing wild-type and embryonic myosin isoforms. Biophys J 98:1218–1226. doi:10.1016/j.bpj.2009.11.051

Elhanany-Tamir H, Yu Y V., Shnayder M, Jain A, Welte M, Volk T. 2012. Organelle positioning in muscles requires cooperation between two KASH proteins and microtubules. J Cell Biol 198:833–846. doi:10.1083/jcb.201204102

Elosegui-Artola A, Andreu I, Beedle AEM, Lezamiz A, Uroz M, Kosmalska AJ, Oria R, Kechagia JZ, Rico-Lastres P, Le Roux AL, Shanahan CM, Trepat X, Navajas D, Garcia-Manyes S, Roca-Cusachs P. 2017. Force Triggers YAP Nuclear Entry by Regulating Transport across Nuclear Pores. Cell 171:1397–1410.e14. doi:10.1016/j.cell.2017.10.008

Fedorchak GR, Kaminski A, Lammerding J. 2014. Cellular mechanosensing: Getting to the nucleus of it all. Prog Biophys Mol Biol 115:76–92. doi:10.1016/j.pbiomolbio.2014.06.009

Forgacs G, Newman SA. 2005. Biological physics of the developing embryo, Biological Physics of the Developing Embryo. doi:10.1017/CBO9780511755576

Fuks NA. 1964. The mechanics of aerosols. Oxford: Pergamon Press.

Gennisson JL, Deffieux T, Macé E, Montaldo G, Fink M, Tanter M. 2010. Viscoelastic and anisotropic mechanical properties of in vivo muscle tissue assessed by supersonic shear imaging. Ultrasound Med Biol 36:789–801. doi:10.1016/j.ultrasmedbio.2010.02.013

Ghannad-Rezaie M, Wang X, Mishra B, Collins C, Chronis N. 2012. Microfluidic chips for in vivo imaging of cellular responses to neural injury in Drosophila larvae. PLoS One 7. doi:10.1371/journal.pone.0029869

Gupta RS. 2015. Elements of Numerical Analysis, second edi. ed. Cambridge University Press.

Hakim CH, Wasala NB, Duan D. 2013. Evaluation of Muscle Function of the Extensor Digitorum Longus Muscle &lt;em&gt;Ex vivo&lt;/em&gt; and Tibialis Anterior Muscle &lt;em&gt;In situ&lt;/em&gt; in Mice. J Vis Exp 1–8. doi:10.3791/50183

Hawley JA, Hargreaves M, Joyner MJ, Zierath JR. 2014. Integrative Biology of Exercise. Cell 159. doi:10.1016/j.cell.2014.10.029

Hector AJ, McGlory C, Phillips SM. 2015. The influence of mechanical loading on skeletal muscle protein turnover. Cell Mol Exerc Physiol 4. doi:10.7457/cmep.v4i1.e8

Hoyt K, Kneezel T, Castaneda B, Parker KJ. 2008. Quantitative sonoelastography for the in vivo assessment of skeletal muscle viscoelasticity. Phys Med Biol 53:4063–4080. doi:10.1088/0031-9155/53/15/004

Janin A, Bauer D, Ratti F, Millat G, Méjat A. 2017. Nuclear envelopathies: A complex LINC between nuclear envelope and pathology. Orphanet J Rare Dis 12:23–42. doi:10.1186/s13023-017-0698-x

Kracklauer MP, Banks SML, Xie X, Wu Y, Fischer JA. 2007. Drosophila klaroid encodes a SUN domain protein required for klarsicht localization to the nuclear envelope and nuclear migration in the eye. Fly (Austin) 1:75–85. doi:10.4161/fly.4254

Lombardi ML, Lammerding J. 2015. HHS Public Access 39:1729–1734. doi:10.1042/BST20110686.Keeping

Martins RP, Finan JD, Farshid G, Lee DA. 2012. Mechanical Regulation of Nuclear Structure and Function. Annu Rev Biomed Eng 14:431–455. doi:10.1146/annurev-bioeng-071910-124638

Méjat A, Misteli T. 2010. LINC complexes in health and disease. Nucleus 1:40–52. doi:10.4161/nucl.1.1.10530

Mellad JA, Warren DT, Shanahan CM. 2011a. Nesprins LINC the nucleus and cytoskeleton. Curr Opin Cell Biol 23:47–54. doi:10.1016/j.ceb.2010.11.006

Mellad JA, Warren DT, Shanahan CM. 2011b. Nesprins LINC the nucleus and cytoskeleton. Curr Opin Cell Biol 23:47–54. doi:10.1016/j.ceb.2010.11.006

Miroshnikova YA, Nava MM, Wickström SA. 2017. Emerging roles of mechanical forces in chromatin regulation. J Cell Sci 130:2243–2250. doi:10.1242/jcs.202192

Mondal S, Ahlawat S, Koushika SP. 2012. Simple Microfluidic Devices for &lt;em&gt;in vivo&lt;/em&gt; Imaging of &lt;em&gt;C. elegans&lt;/em&gt;, &lt;em&gt;Drosophila&lt;/em&gt; and Zebrafish. J Vis Exp 1–9. doi:10.3791/3780

Olatunji O. 2015. Natural Polymers: Industry Techniques and Applications.

Osmanagic-Myers S, Dechat T, Foisner R. 2015. Lamins at the crossroads of mechanosignaling. Genes Dev 29:225–237. doi:10.1101/gad.255968.114

Puckelwartz MJ, Kessler EJ, Kim G, DeWitt MM, Zhang Y, Earley JU, Depreux FFS, Holaska J, Mewborn SK, Pytel P, McNally EM. 2010. Nesprin-1 mutations in human and murine cardiomyopathy Megan. J Mol Cell Cardiol 48:600–608. doi:10.1016/j.yjmcc.2009.11.006

Rajgor D, Shanahan CM. 2013. Nesprins: from the nuclear envelope and beyond. Expert Rev Mol Med 15:e5. doi:10.1017/erm.2013.6

Rindom E, Vissing K. 2016. Mechanosensitive molecular networks involved in transducing resistance exercise-signals into muscle protein accretion. Front Physiol 7:1–9. doi:10.3389/fphys.2016.00547

Rothballer A, Kutay U. 2013. The diverse functional LINCs of the nuclear envelope to the cytoskeleton and chromatin. Chromosoma 122:415–429. doi:10.1007/s00412-013-0417-x

Sloboda DD, Claflin DR, Dowling JJ, Brooks S V. 2013. Force Measurement During Contraction to Assess Muscle Function in Zebrafish Larvae. J Vis Exp 1–10. doi:10.3791/50539

Stroud MJ, Feng W, Zhang J, Veevers J, Fang X, Gerace L, Chen J. 2017. Nesprin 1α2 is essential for mouse postnatal viability and nuclear positioning in skeletal muscle. J Cell Biol 216:1915–1924. doi:10.1083/jcb.201612128

Uhler C, Shivashankar G V. 2017. Regulation of genome organization. Nat Publ Gr 18:717–727. doi:10.1038/nrm.2017.101

Wan LQ, Jiang J, Arnold DE, Guo XE, Lu HH, Mow VC. 2009. NIH Public Access 1:93–102. doi:10.1007/s12195-008-0014-x.Calcium

Wang S, Reuveny A, Volk T. 2015. Nesprin provides elastic properties to muscle nuclei by cooperating with spectraplakin and EB1. J Cell Biol 209:529–538. doi:10.1083/jcb.201408098

Wang S, Stoops E, Unnikannan CP, Markus B, Reuveny A, Ordan E, Volk T. 2018. Mechanotransduction via the LINC complex regulates DNA replication in myonuclei. J Cell Biol 217:2005–2018. doi:10.1083/jcb.201708137

Xie X, Fischer JA. 2008. On the roles of the Drosophila KASH domain proteins Msp-300 and Klarsicht. Fly (Austin) 2:74–81. doi:10.4161/fly.6108

Zhang Q, Ragnauth C, Greener MJ, Shanahan CM, Roberts RG. 2002. The nesprins are giant actin-binding proteins, orthologous to Drosophila melanogaster muscle protein MSP-300. Genomics 80:473–481. doi:10.1016/S0888-7543(02)96859-X

Zhang W, Sobolevski A, Li B, Rao Y, Liu X. 2016. An Automated Force-Controlled Robotic Micromanipulation System for Mechanotransduction Studies of Drosophila Larvae. IEEE Trans Autom Sci Eng 13:789–797. doi:10.1109/TASE.2015.2403393

Zhang Y, Füger P, Hannan SB, Kern J V., Lasky B, Rasse TM. 2010. In vivo Imaging of Intact Drosophila Larvae at Sub-cellular Resolution. J Vis Exp 4–7. doi:10.3791/2249

